# Single-cell RNA sequencing of Tocilizumab-treated peripheral blood mononuclear cells as an in vitro model of inflammation

**DOI:** 10.1101/2020.09.11.281782

**Authors:** Arya Zarinsefat, George Hartoularos, Sindhu Chandran, Chun J. Yee, Flavio Vincenti, Minnie M. Sarwal

## Abstract

COVID-19 has posed a significant threat to global health. Early data has revealed that IL-6, a key regulatory cytokine, plays an important role in the cytokine storm of COVID-19. Multiple trials are therefore looking at the effects of Tocilizumab, an IL-6 receptor antibody that inhibits IL-6 activity, on treatment of COVID-19, with promising findings. As part of a clinical trial looking at the effects of Tocilizumab treatment on kidney transplant recipients with subclinical rejection, we performed single-cell RNA sequencing of comparing stimulated PBMCs before and after Tocilizumab treatment. We leveraged this data to create an in vitro cytokine storm model, to better understand the effects of Tocilizumab in the presence of inflammation. Tocilizumab-treated cells had reduced expression of inflammatory-mediated genes and biologic pathways, particularly amongst monocytes. These results support the hypothesis that Tocilizumab may hinder the cytokine storm of COVID-19, through a demonstration of biologic impact at the single-cell level.

## 1. Introduction

Coronavirus disease 2019 (COVID-19), caused by the severe acute respiratory syndrome coronavirus 2 (SARS-CoV-2), has posed a significant threat to global health since emerging at the end of 2019. Although the spectrum of symptomatic infection ranges significantly, and most infections are not severe^1–3^, the overall global burden of the disease has been significant with up to nearly 20% mortality in certain geographic/demographic groups^4,5^. While notable progress has been made in the understanding the virology and disease process, the abrupt onset and lack of effective vaccination has made treatment of COVID-19 difficult^6,7^.

Interleukin (IL)-6 is a key regulatory cytokine for the innate and adaptive immune response and is a growth factor for B cell proliferation and differentiation, an inducer of antibody production, and a regulator of CD4+ T cell differentiation^8,9^. Early data from the COVID-19 outbreak has shown that the complications from the disease are partly due to increases in various cytokines, including IL-6^10–13^, and that elevated IL-6 levels may be associated with worse outcomes^13–15^. Tocilizumab is an IL-6 receptor antibody, which binds to both the membrane-bound and soluble forms of the IL-6 receptor (IL-6R), thereby inhibiting the action of the cytokine/receptor complex and interfering with the cytokine’s effects^16^. It is a well-studied and accepted therapy for rheumatoid arthritis^17–19^, and has also been studied in giant cell arteritis^20^ and organ transplantation^9,21,22^. As such, multiple global investigators are currently undertaking clinical trials to further assess the efficacy of Tocilizumab in the treatment of COVID-19 and its complications (ClinicalTrials.gov). Thus far, it has been shown that COVID-19 patient plasma inhibits the expression of HLA-DR which may be partially restored by Tocilizumab treatment, and that treatment with Tocilizumab may also improve lymphopenia associated with COVID-19^23^. Preliminary data for Tocilizumab treatment on COVID-19 outcomes has shown improvement in clinical outcomes^24^. While the clinical effects of Tocilizumab in inflammatory and autoimmune disease has been well-studied, there is a paucity of data on the mechanistic/biologic impact of the drug on our immune system.

Given the current state of the COVID-19 epidemic and possible efficacy of IL-6/IL-6R inhibition with the use of Tocilizumab, we believed a deeper analysis of the mechanistic/biologic effects of Tocilizumab could further elucidate the effects of the drug on our immune system. Herein we present an analysis of the impact of Tociluzimab on immune cells using single-cell RNA sequencing (scRNA-seq). We map the response of peripheral blood mononuclear cell (PBMC) subsets to cellular activation using CD3/CD28 stimulation^25–29^. Relevant to understanding the impact of Tociluzimab in suppressing immune activation and inflammation, as seen in the COVID-19 response, we additionally examine the effect of Tociluzimab on unstimulated and stimulated cells, as part of an investigator-initiated clinical trial in kidney transplant (KT) recipients with subclinical rejection (*NIAID U01 AI113362-01*). We provide a resource characterizing the effect of Tociluzimab on immune cells at a single-cell level, and demonstrate the unique and unexpected impact of Tociluzimab on monocytes, and how its effect on suppressing inflammation may be further augmented based on the resting versus activated state of PBMCs before exposing the cells to IL-6R inhibition.

## 2. Results/Discussion

In order to examine the impact of Tocilizumab on the composition and expression of circulating single cells, we compared scRNA-seq data from anti-CD3/CD28 stimulated cells from control (patients not treated with Tocilizumab) PBMCs, to unstimulated PBMCs after 3 to 6 months of Tociluzimab treatment. After filtering cells, a total of 57,737 cells remained for analysis. These cells were put through our analysis pipeline described (see Methods). After UMAP clustering, there were a total of 21 distinct clusters representing major PBMC groups, inclusive of naïve CD4+ T/CD8+ T, activated CD4+ T/CD8+ T, memory CD4+ T/CD8+ T, B, Natural Killer (NK, both CD56+ dim and bright cells^39^), dendritic (DC) cells, and monocytes. Clusters were annotated according to canonical cell type markers (*<u>Figure 1A</u>*). Clusters 2, 4, and 5 expressed markers of memory T cell expansion (*S100A4, IL7R*^40,41^) while clusters 0, 8, and 16 expressed markers of CD4+ T cell activation (*TNFRS4, CD69*^42^). One cluster of doublets (cluster 20) was removed to give the final annotated clusters (*<u>Figure 1B</u>*).

**Figure 1:**
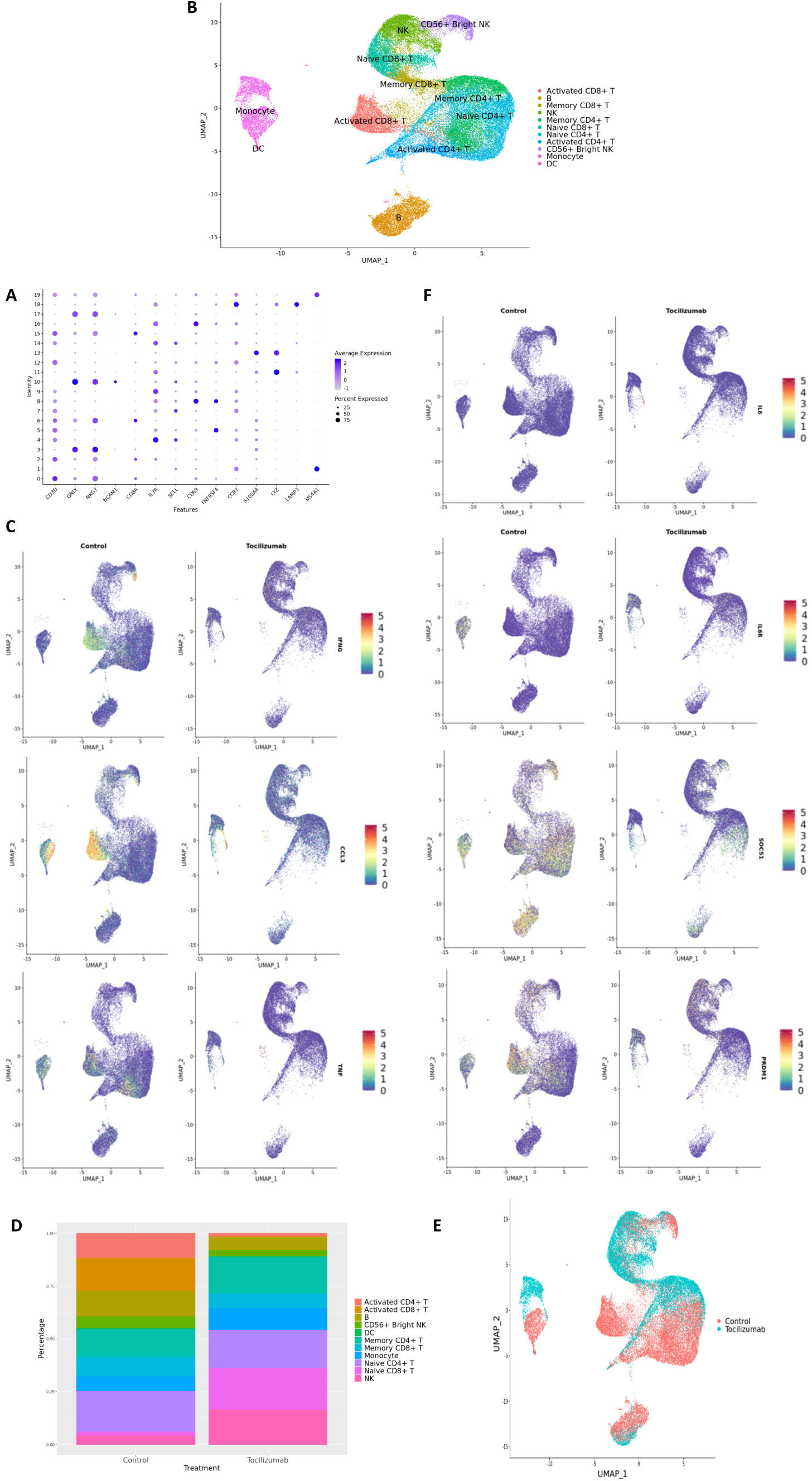
UMAP clustering, cell subset annotation, and expression of inflammatory markers and IL6R pathway genes in control vs. Tocilizumab-treated PBMCs **a**, Dot plot of canonical markers used for annotation of the 20 cell clusters. Average feature expression represented by color gradient with lower expression represented by light grey, and higher expression represented by blue. Size of dots represent the percent of cells within that specific cluster that express the feature of interest **b**, UMAP with final cell type annotations **c**, Feature plots showing expression of select cytokines involved in SARS-CoV-2 cytokine storm (*IFNG, CCL3*, and *TNF*) based on control vs. Tocilizumab treatment status. Feature expression represented by color gradient, with low expression represented by blue and high expression represented by red **d**, Bar plot showing the percentage of each cell type in control vs. Tocilizumab-treated groups **e**, UMAP with cell clusters identified based on control vs. Tocilizumab treatment status **f**, Feature plots showing expression of *IL6, IL6R*, and downstream IL6R pathway genes (*SOCS1, PRDM1*) based on control vs. Tocilizumab treatment status. Feature expression represented by color gradient, with low expression represented by blue and high expression represented by red

Feature plots showing the expression of “cytokine storm”^43^related pro-inflammatory genes are cell-type specific, with predominance for expression in T cell and monocyte clusters (*<u>Figure 1C</u>*). Although many genes are known to be involved in the cytokine storm of COVID-19^37,38^, we demonstrate that some of the key pro-inflammatory genes (cytokines, interferons, and tumor necrosis factor) are also noted as part of the inflammatory profile in control (no Tocilizumab) patients (*<u>Figure 1C</u>*, control cells). Overall, stimulated PBMCs not exposed to Tociluzimab show a dominant signal for T cell activation. After 6 months of treatment with Tociluzimab there is a shift in peripheral blood subset frequencies observed across no treatment (control) vs. treatment (Tocilizumab) groups. In comparison to changes in overall cell types, there was little observed effect on frequencies of naïve CD4+/CD8+ T cells, DC, or NK cells, but with a marked reduction of activated CD4+ T cells (approximately 12.5% of control PBMCs were activated CD4+ T cells, while there were essentially no activated CD4+ T cells in the Tocilizumab group, *<u>Figure 1D</u>*). Within these different cell subsets, Tociluzimab therapy results in significant polarization of gene expression based on UMAP presentation (*<u>Figure 1E</u>*), with notable polarization by treatment status observed in monocytes.

Given Tocilizumab’s function as an IL-6R blocker, we looked at the expression of *IL6, IL6R*, as well as *SOCS1* (feedback inhibitor of IL-6 signaling, expressed upon IL-6 pathway activation^44^), and *PRDM1* (activated by the *JAK/STAT3* pathway via activation of the IL-6 pathway ^45,46^) in Tocilizumab-treated cells (*<u>Figure 1F</u>*). Tocilizumab-treatment resulted in the expected reduction of *IL6R, SOCS1* and *PRDM1* expression, in CD4+ and CD8+ T cells, and unexpectedly also in monocytes. *IL6* expression did not appear to be affected by Tocilizumab treatment.

We then looked at the top 30 most differentially expressed genes (highest log_2_-fold changes) for control vs. Tocilizumab amongst all cells (*<u>Figure 2A</u>*), CD4+ T cells (*<u>Figure 2B</u>*), CD8+ T cells (*<u>Figure 2C</u>*), monocytes (*<u>Figure 2D</u>*), and performed corresponding PA for these genes. PA showed enrichment of inflammatory pathways such IL and TNF signaling amongst control cells. We looked at the top 30 most differentially expressed genes (highest log_2_-fold changes) for control vs. Tocilizumab monocytes (*<u>Figure 2D</u>*), with some notable differences as would be expected. Control monocytes were enriched in chemokines such as *CXCL9*, various HLA genes involved in antigen processing^48^ (*HLA-DQB1, HLA-DRB5*), *CD40* (member of the TNF-receptor superfamily^49^), and *SOCS1* (downstream gene activated by IL-6R pathway, as previously discussed^44^). PA revealed enrichment of many inflammation-related pathways, including interferon, interleukin, T cell receptor (TCR), and PD-1 signaling in control PBMCs, suggesting the relative suppression of these pathways in cells exposed to Tocilizumab.

**Figure 2:**
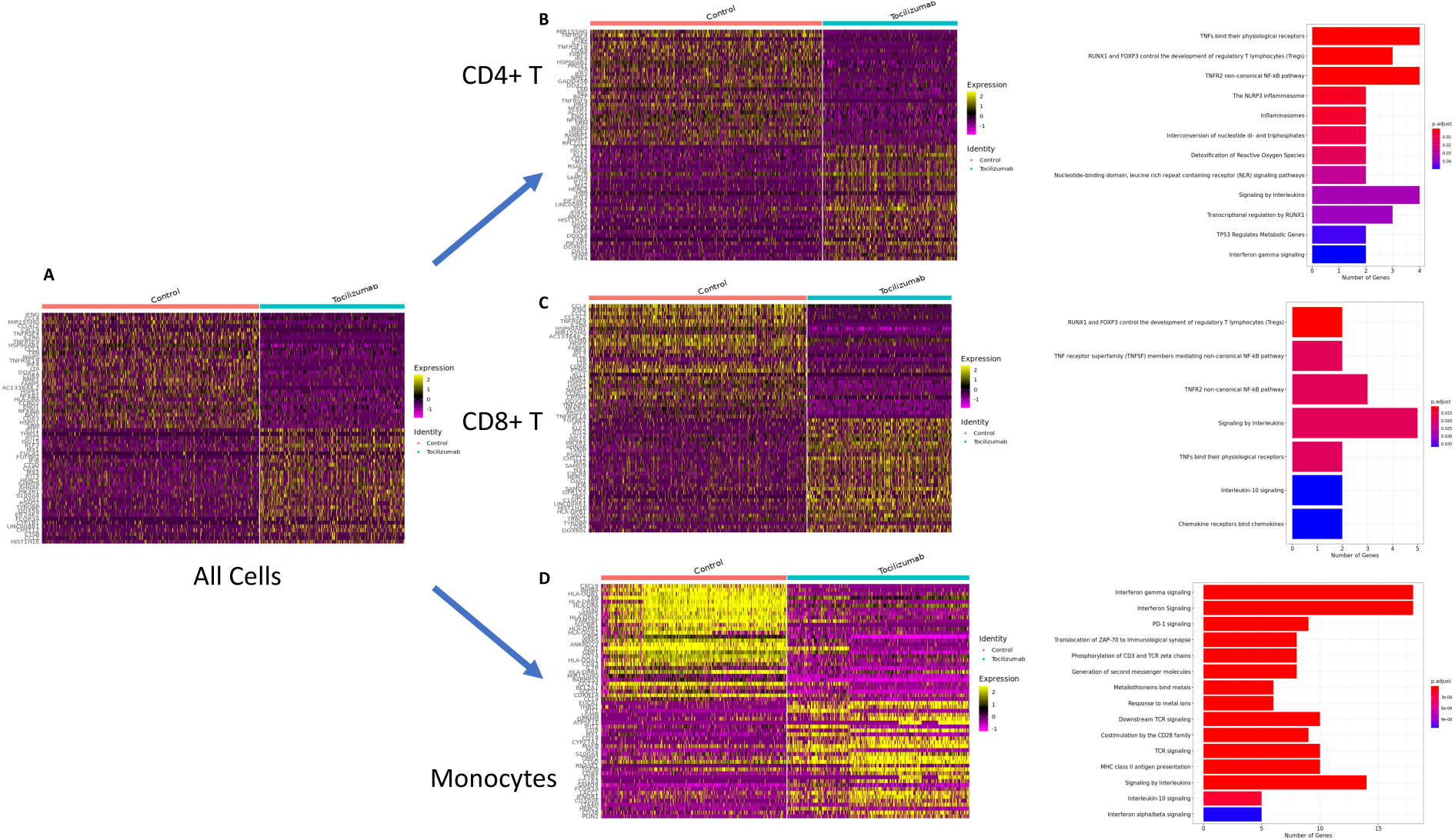
Differential expression testing and pathway analysis of all cells, CD4+ T cells, CD8+ T cells, and monocytes **a**, Heatmap of top 30 genes with highest log-fold changes in all control and Tocilizumab-treated cells **b**, Heatmap of top 30 genes with highest log-fold changes in all CD4+ T control and Tocilizumab-treated cells, with corresponding PA of top 10% most highly differentially expressed genes (based on log_2_-fold change) in control vs. Tocilizumab cells **c**, Heatmap of top 30 genes with highest log-fold changes in all CD8+ T control and Tocilizumab-treated cells, with corresponding PA of top 10% most highly differentially expressed genes (based on log_2_-fold change) in control vs. Tocilizumab cells **d**, Heatmap of top 30 genes with highest log-fold changes in all control and Tocilizumab-treated monocytes, with corresponding PA of top 10% most highly differentially expressed genes (based on log_2_-fold change) in control vs. Tocilizumab cells. Gene expression level represented by color gradient ranging from purple (low expression) to yellow (high expression). PA figure x-axis represents the number of genes from each pathway that was present in the gene list. Adjusted p-values for pathway enrichment are represented as a color gradient with larger p-values colored blue and smaller p-values colored red

In addition to the effect of Tociluzimab on T cells, we also observed an unexpected polarization of monocytes after Tocilizumab treatment (*<u>Figure 1E</u>*). Notably, the Tocilizumab monocyte cluster was enriched for *CD14*, suggestive of an increased presence of classical monocytes^47^, while *CD16*/*FCGR3A* expression was more evenly expressed between the two clusters (*<u>Figure 3A</u>*). We then performed cell trajectory analysis of these monocytes for Tociluzimab treatment effect, utilizing *Monocle*. This revealed six distinct cell trajectory branches, with two of the branches containing nearly all control cells not exposed to Tocilizumab, and the other four branches containing nearly all Tocilizumab-exposed PBMCs (*<u>Figure 3B</u>*), supporting the presence of unique PBMC trajectories after patient exposure to IL6-R blockade. We utilized *Monocle’s* BEAM function to perform branched expression analysis modeling of the distinct cell trajectory branches for Tociluzimab-exposed PBMCs (circled branch, *<u>Figure 3B</u>*), which showed distinct clusters of cells based on treatment status (*<u>Figure 3C</u>*).

**Figure 3:**
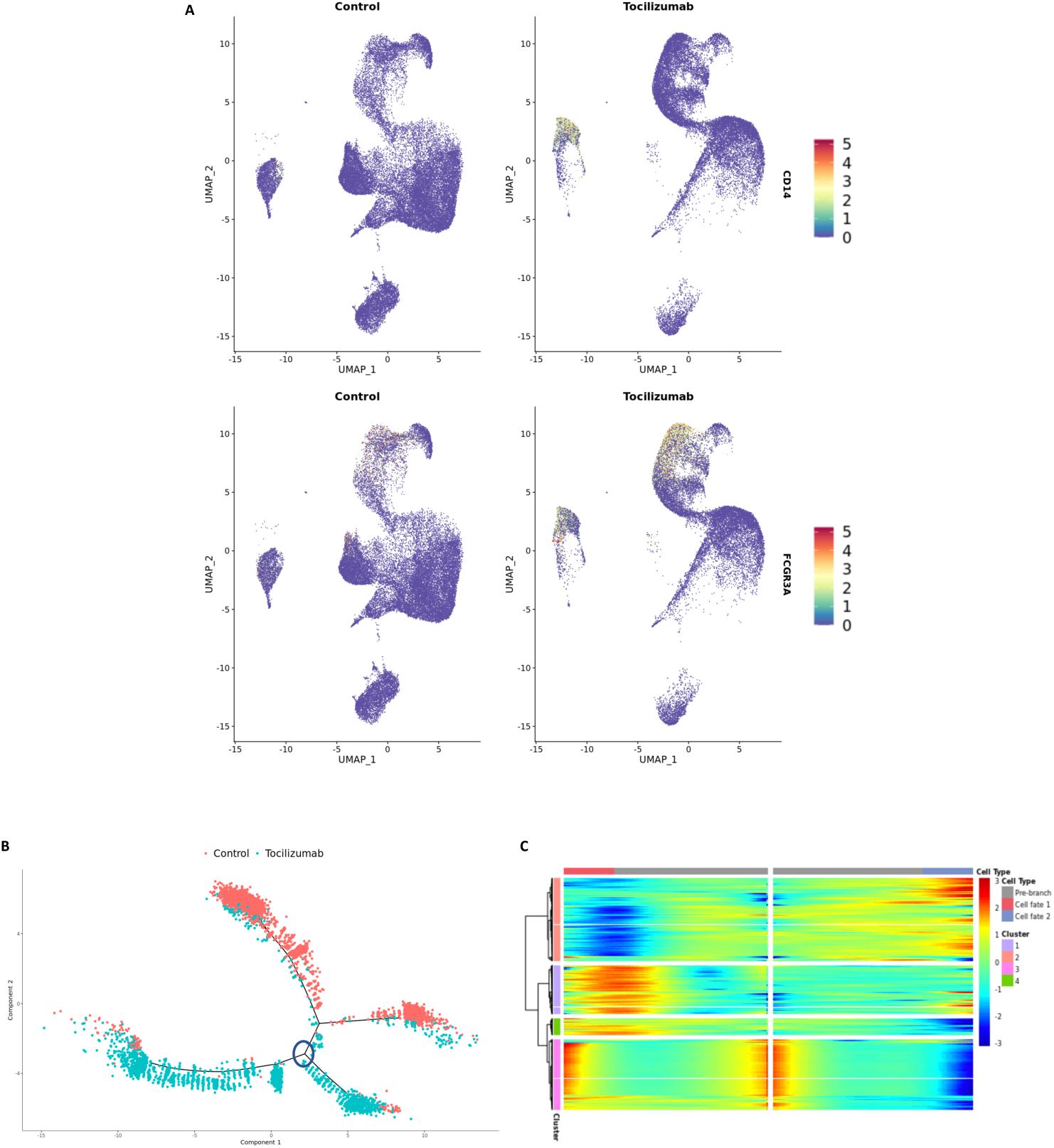
Differential expression testing and cell trajectory analysis of monocyte subsets **a**, Feature plots showing expression of CD14 and CD16 based on control vs. Tocilizumab treatment status. Feature expression represented by color gradient, with low expression represented by blue and high expression represented by red, with higher CD14 expression noted in Tocilizumab cells **b**, Cell trajectory analysis of monocyte clusters showing distinct lineages of control vs. Tocilizumab cells; blue circle represents branch point used in subsequent heatmap analysis **c**, Heatmap from branched expression analysis modeling for most differentially expressed genes between branch points from **b** (analyzed branch point marked by blue circle), showing clusters of differentially expressed genes between branches. Gene expression represented as color gradient from blue (low expression) to red (high expression). Cell type annotation represented by two separate cell fates as seen in **b**, with middle of heatmap representing the start of pseudotime and clear separation of control vs. Tocilizumab cell fates

The results of this study showed that in PBMCs undergoing a cytokine storm signal in rejection^50^, with overlapping signatures of *IFNG, CCL3*, and *TNF* expression, along with TCR signaling also seen in the cytokine storm of COVID-19^37,38^, there is suppression of these inflammatory pathways after Tocilizumab treatment. This is inclusive of suppression of downstream signaling of IL6-R pathway genes in both monocytes and T cells.

Monocytes have been shown to play a significant role in the pathophysiology of COVID-19^51^. A significant expansion of populations of monocytes producing IL-6 has been observed in the peripheral blood of patients with COVID-19 in ICUs compared with those patients who did not require ICU hospitalization^52^, with similar findings of increased IL-6 production from monocytes also seen by scRNA-seq analysis of PBMCs^53^. Our findings are from the first clinical trial utilizing Tocilizumab for transplant rejection recipients and the first scRNA-seq analysis for such a study. We show a separation of cell clustering based on treatment status, reduced enrichment of inflammatory pathways in Tocilizumab patients, and relatively reduced expression of IL-6R pathway genes in Tocilizumab-treated cells. As would be expected, we did not observe any differences in IL-6 gene expression between control and Tocilizumab cells (as Tocilizumab is an IL-6R blocker), but rather only effects on the subsequent function of that cytokine’s pathways. We also show an enrichment of *CD14* expression (associated with classical monocytes) in Tocilizumab-treated monocytes, which are believed to be phagocytic, but with reduced inflammatory attributes^47^. This is consistent with our PA described above that shows enrichment of inflammatory pathways in control cells, but not Tocilizumab-treated cells (possibly due to the increased presence of non-inflammatory classical monocytes in Tocilizumab-treated cells).

Our findings, in conjunction with the available data on clinical outcomes of Tocilizumab treatment^24^ and ongoing trials, show promise for the use of Tocilizumab in the treatment of patients with COVID-19. The results of our study support the belief that Tocilizumab may be effective in reducing the inflammatory burden that results in the adverse outcomes of COVID-19. Future studies will need to be undertaken to look at outcomes of Tocilizumab treatment for COVID-19 in a clinical trial setting, ideally in conjunction with scRNA-seq analysis of these patient’s blood samples to achieve a greater understanding of the transcriptomic effects of infection and treatment at a single-cell level.

## 3. Materials and Methods

### Sample collection

This study was performed as part of an ancillary to a randomized controlled clinical trial of 15 KT recipients that were diagnosed with subclinical rejection on their 6-month post-transplant protocol biopsy and randomized to either continue standard of care (Tacrolimus, mycophenolate, and steroid) immunosuppression (control arm, 8 patients) or standard of care plus Tocilizumab (Tocilizumab treatment arm, 7 patients). Patients in the treatment arm were given Tocilizumab at a dose of 8 mg/kg IV every 4 weeks, for a total of 6 doses. Patients in both arms of the study had blood collected at baseline prior to the initiation of Tocilizumab (in the treatment arm patients), then at 3, 6, and 12 months after the start of the study, for a total of 4 blood samples per all 15 patients in the study. PBMCs were isolated from blood samples by Ficoll-Paque™ PLUS density gradient centrifugation (GE Healthcare, Chicago, IL) and frozen in fetal bovine serum (Gibco, Waltham, MA) containing 10% (vol/vol) dimethyl sulfoxide (Sigma-Aldrich, St. Louis, MS). Cells were frozen and not thawed until the day of the experiment when they were used directly for in vitro stimulation.

### Stimulation with anti-CD3 and anti-CD28 antibodies

Frozen PBMCs were thawed, four vials at a time to ensure maximum cell recovery, in a water bath at 37 Celsius. Cells were counted using a hematocytometer, split in half, and were then adjusted to 2×10^5^ cells/well and triplicate plated in multiscreen 96-well plates (Falcon, Corning, NY). Cells were stimulated with soluble anti-CD3 (5 μg/mL; MabTech, Cincinnati, OH) and anti-CD28 antibodies (10ug/mL; MabTech, Cincinnati, OH) at 37 Celsius, 5% CO_2_ for 24 hours. Unstimulated PBMCs were incubated under identical conditions to reduce any confounding from incubation conditions other than stimulation. Since all PBMCs were split in half prior to any downstream processing, all samples from control and Tocilizumab-treated patients at all study time points were both stimulated and not stimulated as part of the study design.

### Sample processing

After overnight stimulation/incubation, the cells were harvested and counted using a hematocytometer and orange acridine solution. Any cell suspension that was less than 25 cells/uL was disqualified from multiplexing due to low cell counts. Multiplexing cell pools were designed such that no pair of stimulated and unstimulated samples from the same patient were in the same pool and such that no samples from the same collection time point were in the same pool. The same number of cells from each patient and experimental condition were multiplexed into their respective pools to make a final total of 300,000 cells per pool. Any remaining non-pooled cells were resuspended in RNAlater (Thermo-Fisher, West Sacramento, CA) and saved for SNP array. Cell pools were then centrifuged at 400g for 5 minutes and media was aspirated. Cell pellet was resuspended in a small volume of Wash Buffer (0.4% BSA in 1XPBS) and the suspension was filtered through a 40uM cell strainer (Falcon, Corning, NY).

### Library construction and sequencing

scRNA-seq libraries were prepared using the 10X Chromium Single Cell 3’ Reagent Kits v3, according to the manufacturer’s instructions. Briefly, the isolated cells were washed once with PBS + 0.04% BSA and resuspended in PBS + 0.04% BSA to a final cell concentration of 1000 cells/μL as determined by hematocytometer. Cells were captured in droplets at a targeted cell recovery of 4000-8000 cells, resulting in estimated multiplet rates of 0.4-5.4%. Following reverse transcription and cell barcoding in droplets, emulsions were broken and cDNA purified using Dynabeads MyOne SILANE (Thermo-Fisher, West Sacramento, CA) followed by PCR amplification (98°C for 3 min; 12-16 cycles of 98°C for 15 sec, 67°C for 20 sec, 72°C for 1 min; 72°C for 1 min). Amplified cDNA was then used for 3’ gene expression library construction. For gene expression library construction, 2.4-50 ng of amplified cDNA was fragmented and end-repaired, double-sided size selected with SPRIselect beads (Beckman Coulter, West Sacramento, CA), PCR amplified with sample indexing primers (98°C for 45 sec; 14-16 cycles of 98°C for 20 sec, 54°C for 30 sec, 72°C for 20 sec; 72°C for 1 min), and double-sided size selected with SPRIselect beads. Pooled cells were loaded in a 10X chip in three replicate wells such that each well contained 50,000 cells. Given the large number of cells and large number of patient samples, the entire experiment and sequencing was performed in 2 separate batches to prevent cell death during counting. Each day resulted in 4 unique pools with each pool run in triplicate wells for sequencing. Sequencing single-cell RNA libraries were sequenced on an Illumina NovaSeq S2 to a minimum sequencing depth of 50,000 reads/cell using the read lengths 26bp Read1, 8bp i7 Index, 91bp Read2.

### Demultiplexing

To assign cells to donors of origin in our multiplexed design, we leveraged the genetic demultiplexing tools *demuxlet*^30^ and *freemuxlet*, both a part of the *popscle* suite of population genetics tools (https://github.com/statgen/popscle). These tools leverage the genetic polymorphisms present in transcripts to assign the cells found in each droplet to their donor of origin. *Demuxlet* uses the genotype calls from a genotyping SNP array to classify cells in droplets according to their donor of origin, while freemuxlet “learns” the genotypes of a pre-defined number of donors from the transcripts themselves, and assigns the droplets to a respective anonymous donor according to those learned genotypes. Upon first receiving sequencing data, *demuxlet* was run with input genotypes from all the patients in the cohort. While *demuxlet* was able to assign most droplets to donors of origin, it revealed that two patients in the genotyping SNP array appeared to have identical genotypes (likely due to human error) and that cells from some patients seemed to drop out (likely due to low viability cells or inaccurate cell counting or mixing). Therefore, to validate *demuxlet* results, *freemuxlet* was run using an independent list of SNP sites: exonic SNPs with a minor allele frequency > 0.05 as observed in the 1000 Genomes Project. In order to leverage the droplets across multiple microfluidic reactions, which may enable higher confidence in the learned genotypes, we merged the BAMs from multiple experiments containing the same patients into a single BAM and input this merged BAM into *freemuxlet*. The droplet assignments from the anonymous donors output by *freemuxlet* were then compared to those from *demuxlet*, showing very high concordance. Moreover, comparing the VCF generated from *freemuxlet* (using the SNPs present in the droplets) to the VCF generated from the SNP genotyping array yielded a 1:1 correspondence of anonymous individuals to patients, barring those few problematic patients. Through comparing VCFs and the presence/absence of individuals in each multiplexed experiment, we were able to definitively assign a detected genotype to all detected individuals. Droplet barcodes were then filtered to remove heterotypic droplets containing cells from multiple individuals, and the remaining homotypic droplets were analyzed downstream.

### Data analysis

Raw FASTQ files were processed using *CellRanger* (v 3.0.1) to map reads against human genome 38 as a reference, filter out unexpressed genes, and count barcodes and unique molecular identifiers (UMIs). Subsequent analyses were conducted with *Seurat* (v 3.1.2)^31^ in *R* (v 3.6.2). We compared PBMCs from all anti-CD3/CD28 stimulated cells from the study baseline, to unstimulated Tocilizumab-treated cells from 3 to 6 months post-treatment with Tocilizumab. Utilizing *Seurat*, we first filtered cells to only keep those that had less than 10% mitochondrial genes and cells with numbers of features greater than 200 and less than 2,500. Cells were assigned patient identification based on the *demuxlet/freemuxlet* output described above, and once patients were identified, additional treatment/stimulation/time metadata could be applied. Given that our experiment was divided over 2 days given the high number of samples/cells, we applied *Seurat’s* SCTransform function for data integration to account for any possible batch effects from experiment days^32,33^. Once the data was integrated, we continued downstream data processing. We first determined the principal components (PCA), then constructed a shared nearest neighbor graph (SNN), identified clusters with a resolution of 0.75, and finally visualized the cells using uniform manifold approximate and projection (UMAP), per the typical Seurat workflow^31^. Clustering was achieved by using 15 components from the PCA dimensionality reduction.

To identify cluster-specific markers following the creation of UMAP plots, we utilized normalized RNA counts of all clusters, scaled the data, and performed differential gene expression (DE) testing by applying the Wilcoxon rank sum test using *Seurat’s* FindMarkers function^31^. We also plotted normalized and scaled gene expression of canonical markers in conjunction with DE testing to determine identities of each cluster. To compare cell clusters of stimulated vs. unstimulated cells, or control vs. Tocilizumab-treated cells, we once again utilized normalized/scaled RNA counts and performed DE testing with FindMarkers.

To perform pathway analysis (PA) for any specific comparison we performed, we filtered for all differentially expressed genes with an adjusted (based on the Bonferroni correction) p-value < 0.05, and then selected the top 10 percentile of genes with the highest log-fold changes. These top genes were used to perform the PA utilizing the Reactome database^34^ with the *clusterProfiler* package^35^. To perform cell trajectory analysis, we first subset our clusters and cell types of interest from our *Seurat* workflow, then performed dimensionality reduction and cell ordering with *Monocle*^36^ (v 2.14.0). We were then able to plot specific cells by their trajectory branches based on their pseudotime values assigned by *Monocle*. DE of individual cell trajectory branches was then performed with *Monocle’s* BEAM (branched expression analysis modeling) function, followed by visualization of these differentially expressed branches with *Monocle’s* heatmap visualization tool.

## Acknowledgements

The authors thank the many individuals without whose enthusiastic participation and help this study would never have been accomplished. We would like to acknowledge the following: TA Sigdel who contributed to the study design, JA Liberto who contributed to study design and cell culture/isolation, P Rashmi who contributed to the study design, and AA Da Silva who contributed to cell culture/isolation. A Zarinsefat is funded by the NIH: 5 T32 AI 125222.

